# Evaluating cytotoxicity and genotoxicity of oil extracted from visceral fat of *Caiman yacare* (Daudin, 1802) in chinese hamster lung fibroblast *in vitro*

**DOI:** 10.1101/2023.07.28.551009

**Authors:** Lucas P. Azevedo, Fabricio Rios-Santos, Carmen L. B. Branco, Leandro N. Pressinotti, Érica de M. Reis, Samuel V. Filho, Domingos T. de O. Martins, Willian de Arruda Silva, Leonardo G. de Vasconcelos, Rosa Helena dos Santos Ferraz, Fernanda Vieira Mesquita, Paulo T. S. Junior

## Abstract

In previous studies, the oil extracted from the visceral fat of *Caiman yacare* (Daudin, 1802) demonstrated a wound-healing effect on the skin of Wistar rats. To enhance knowledge our about the mechanism underlying this effect, we analysed the oil’s toxicological potential *in vitro*. Cytotoxicity, genotoxicity, pro-oxidant, and antioxidant activities were evaluated in a V79-4 cell line. The oil was obtained using the Soxhlet method, and the proportions of the fatty acid profile was previously identified 43.74 % saturated and 34.65 % unsaturated fatty acids. Protocol 487 of the Organisation for Economic Co-operation and Development (OECD) was employed for cell line selection and concentrations. Cytotoxicity was determined using the MTT assay and clonogenic survival. Pro-oxidant and antioxidant activities were analysed using flow cytometry. Genotoxicity was evaluated using comet and micronucleus assays. The oil did not demonstrate cytotoxicity up to a concentration of 500 µg/mL. At concentrations of 250 and 500 µg/mL, the oil exerted a protective effect against oxidative stress and showed genotoxic effects only at the highest concentration (2000 µg/mL). Like other oils of interest for human health, the oil extracted from the visceral fat of *C. yacare* demonstrated low toxicological potential *in vitro*.

**SUMMARY STATEMENT:** The oil from *Caiman yacare* visceral fat presents low cytotoxicity and genotoxicity, highlighting its potential for therapeutic applications without adverse effects.

## 1. INTRODUCTION

*Caiman yacare* is a reptilian species found in the Brazilian Pantanal, with high economic value owing to its meat and leather (Vicente-Neto et al., 2010). However, the growing awareness of sustainability issues has led consumers to seek products with the lowest environmental impact (Kumar et al., 2021). Captive management of *C. yacare* using the ranching method meets this requirement by monitoring the area and conservation of the local biome (Campos et al., 2020).

However, approximately 2.65 kg of viscera are discarded per animal during their slaughter (Romanelli and Schmidt, 2003). This wasted viscera include tissues and organs rich in fats, which are used in the production of cosmeceuticals and nutraceuticals (Shim-Prydon and Camacho-Barreto, 2007). In Brazil, the northeastern and traditional communities in the Amazon, especially indigenous people, frequently use crocodilian oils and fats as a food source and for the treatment of diseases (Ferreira Abrao et al., 2021; Mishra et al., 2020). Crocodilian fats and oils are used in the treatment of wounds and infectious diseases, and as an anti-inflammatory agent (Alves et al., 2017).

The wound healing property of oil extracted from visceral fat of *C. yacare* (hereafter referred to as ‘CVO’) and *Crocodylus siamensis* was demonstrated in a rat excisional wound model (Azevedo et al., 2020) and in rat burns, respectively (Li et al., 2021). Evidence has indicated the functional and chemical similarity between oils from different crocodilian species, but safety data is only available for *C. siamensis* oil. The oil from this species showed no toxicity in orally supplemented rats (Praduptong et al., 2018), did not decrease the cell viability of macrophages, and demonstrated a protective effect against damage caused by oxidative stress on macrophage DNA (Ngernjan et al., 2022).

*In vitro* cytotoxic and genotoxic assays are required by regulatory agencies to determine an effective and safe therapeutic range, before animal testing can be conducted (Kramer et al., 2007). Therefore, due to the central role played by DNA in the homeostasis of living organisms, an *in vitro* cytotoxic and genotoxic evaluation becomes indispensable in the development of wound healing products.

In this study, the *in vitro* cytotoxicity and genotoxicity of oil extracted from visceral fat of *Caiman yacare* (CVO) were investigated, using the V79-4 cell line of Chinese hamster lung fibroblasts.

## 2. RESULTS

### 2.1 Cytotoxicity: cell viability and clonogenic survival assay

CVO did not reduce cell viability at any of the evaluated concentrations [*f* (9, 20) = 7.97, *p* ≥ 0.05]. In contrast, the CVO increased the conversion of MTT salt into formazan blue at concentrations up to 1000 µg/mL (Fig. 1A). Cells treated with CVO may be metabolically active, but with compromised proliferative capacity due to DNA damage; however, the MTT assay cannot measure this effect. Therefore, the clonogenic survival assay was performed to verify the cytotoxic effects of CVO on both proliferation and cell death. The results demonstrate that CVO treatment reduced the SF only at the two highest concentrations (1000 and 2000 µg/mL) [*f* (5, 12) = 22.26, *p* ≤ 0.0001] (Fig. 1B). The cytostatic effect, another parameter of cytotoxicity of CVO treatments, was below 30% [*f* (5, 12) = 26.38, *p* ≤ 0.05], confirming the results observed in the clonogenic survival assay (Fig. 1C).

**Fig. 1.**
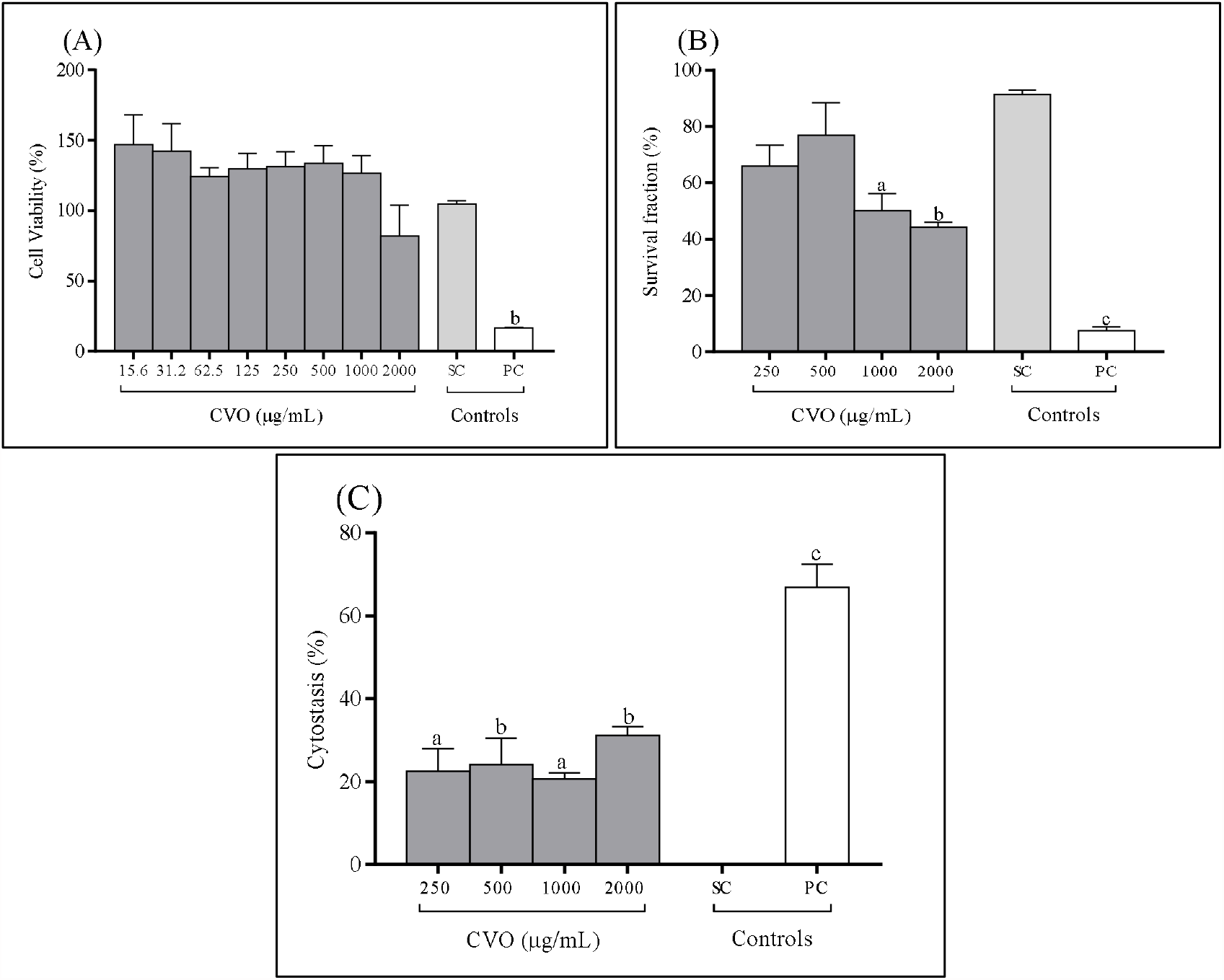
Cytotoxicity of Chinese hamster lung fibroblast, V79-4 cell line, after treatment with different concentrations of oil extracted from visceral fat of *Caiman yacare* (CVO) for 24 h. (A) Cell viability (MTT assay); (B) survival fraction (SF, clonogenic survival assay); (C) amount of cytostasis (%). Values refer to mean ± standard error of the mean (SEM) of three independent assays in triplicate. One-way ANOVA test followed by Dunnett’s post-hoc test. Letters above the bars indicate significance; a, p ≤ 0.05; b, p ≤ 0.01; and c, p ≤ 0.001 versus the solvent control (SC, 0.08% DMSO). Positive control (PC, doxorubicin 50 mM, MTT assay) and (PC, doxorubicin 0.025 µg/mL, clonogenic survival assay, cytostasis).

### 2.2 Determining intracellular ROS levels

The treatment of cells with CVO for 4 h resulted in a significant increase in ROS levels (pro-oxidant) only at the highest concentration (2000 µg/mL) [*f* (5, 12) = 19.6, *p* ≤ 0.0001] compared to positive control (PC, H_2_O_2_) (Fig. 2A). CVO also demonstrated an antioxidant effect, reducing ROS levels in cells treated with concentrations of 250 and 500 µg/mL [*f* (6, 14) = 3.43, *p* = 0.026] when compared to positive control (PC, NAC) (Fig. 2B).

**Fig. 2.**
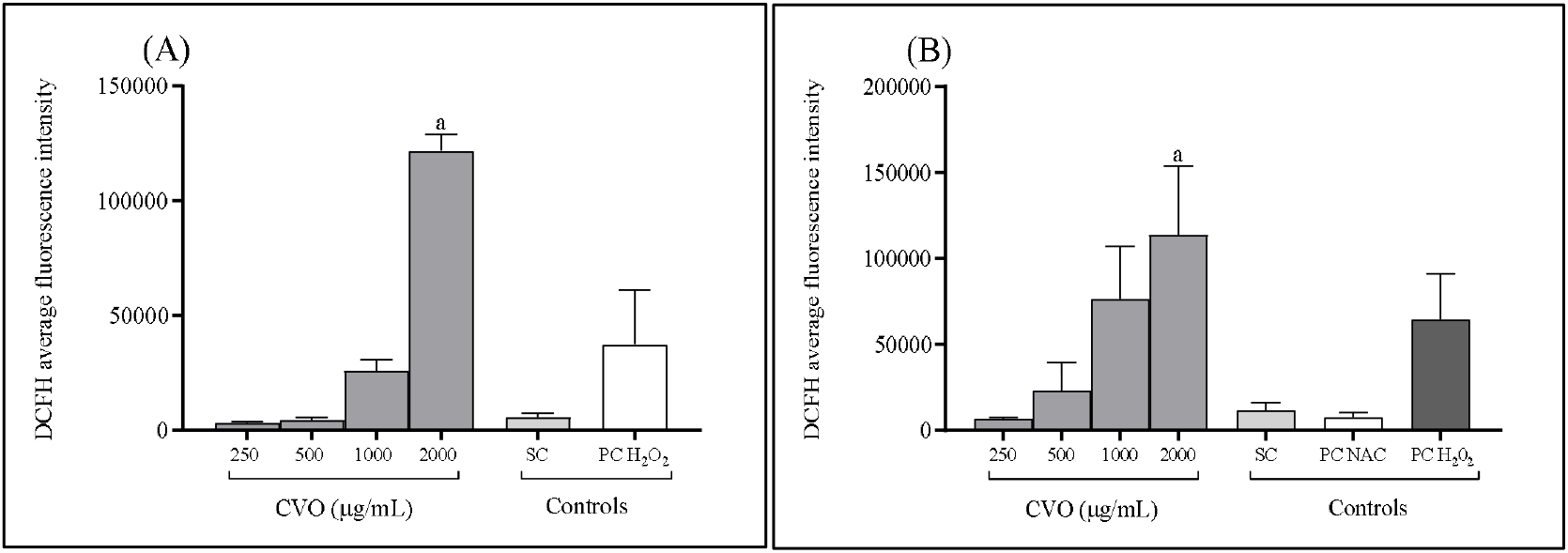
Intracellular ROS production in Chinese hamster lung fibroblasts, V79-4 cell line, treated with different concentrations of oil extracted from visceral fat of *Caiman yacare* (CVO). (A) For pro-oxidant activity (ROS production), the cells were exposed to different concentrations for 4 h. (B) For antioxidant activity (protective effect), cells were treated for 3 h. Then, treatment solutions were removed, and oxidative stress was induced with hydrogen peroxide (H2O2) at 1 mM for 1 h in all cultures. Data are presented as mean ± SEM of three independent assays. One-way ANOVA test, followed by Dunnett’s post-hoc test. Letters above the bars indicate significance; a, p ≤ 0.05 versus hydrogen peroxide positive control (PC, H2O2; 1 mM, pro-oxidant) and positive control with N-acetyl cysteine (PC, NAC; 2 mM, antioxidant).

### 2.3 Evaluation of genotoxicity

#### 2.3.1 Comet assay

CVO did not increase the percentage of single-strand DNA breaks [*f* (5, 12) = 5.40, *p* ≥ 0.05] (Fig. 3A). However, it increased the percentage of oxidative DNA damage, but only at the highest concentrations tested (1000– 2000 µg/mL) [*f* (5, 12) = 11.44, *p* = 0.0003] (Fig. 3B). Interestingly, treatment with the lowest concentration (250 µg/mL) reduced the percentage of oxidative damage compared to the solvent control.

**Fig. 3.**
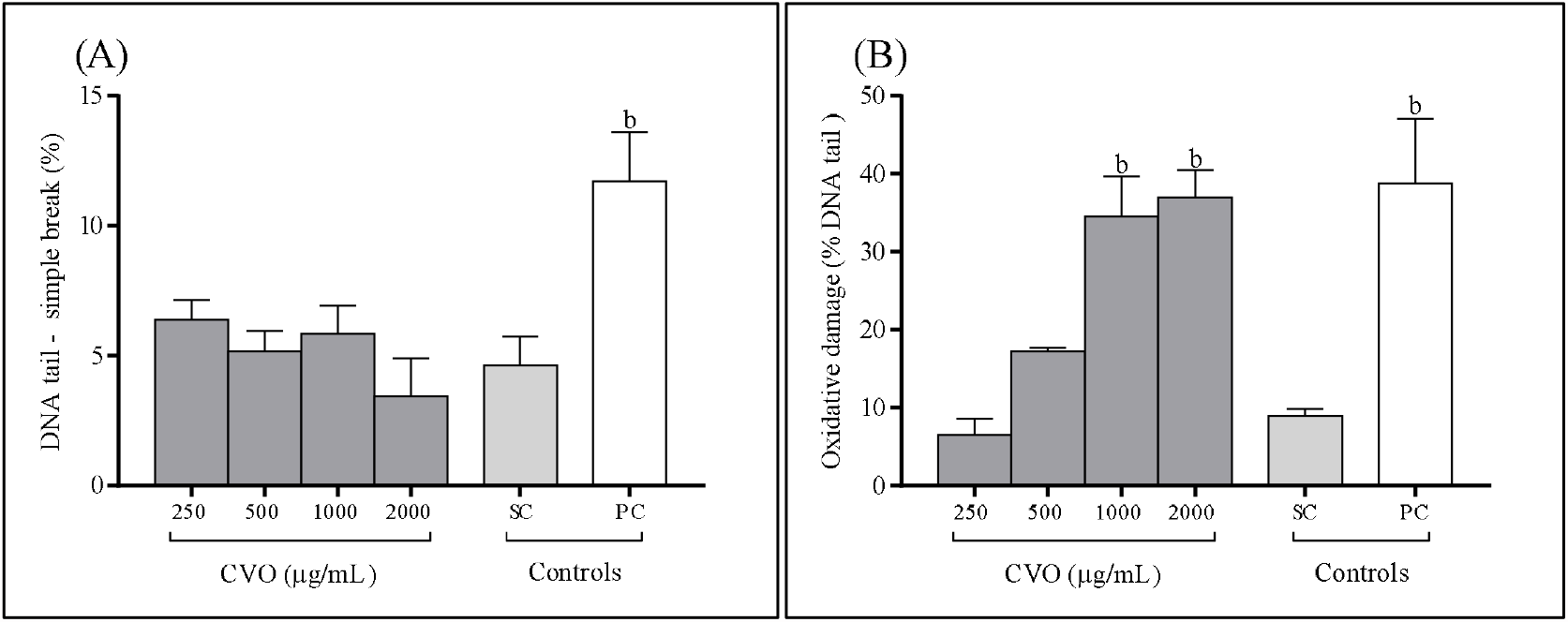
Percentage of DNA damage in Chinese hamster lung fibroblasts, V79-4 cell line, after treatment with different concentrations of oil extracted from visceral fat of *Caiman yacare* (CVO) for 3 h. (A) Single-strand breaks in DNA (standard alkaline comet assay); (B) oxidative DNA damage (modified alkaline comet assay with the repair enzyme formamidopyrimidine(fapy)-DNA glycosylase (Fpg). Data are presented as the mean ± SEM of three independent assays. One-way ANOVA test, followed by Dunnett’s post-hoc test. Letters above the bars indicate significance; a, p ≤ 0.01 versus solvent control (SC, 0.8% DMSO). Positive control (PC, doxorubicin 0.025 µg/mL).

#### 2.3.2 Cytokinesis-block micronucleus assay (CBMN)

Cells treated with CVO had increased numbers of micronuclei (MN) [*f* (5, 12) = 5.22, *p* = 0.008], nuclear buds (Nbuds) [*f* (5, 12) = 27.77 *p* ≤ 0.0001] and nucleoplasmic bridges (NPB) [*f* (5, 12) = 12.81, *p* ≤ 0.0002] only at higher concentrations (1000–2000 µg/mL) (Fig. 4A, B, and C).

**Fig. 4.**
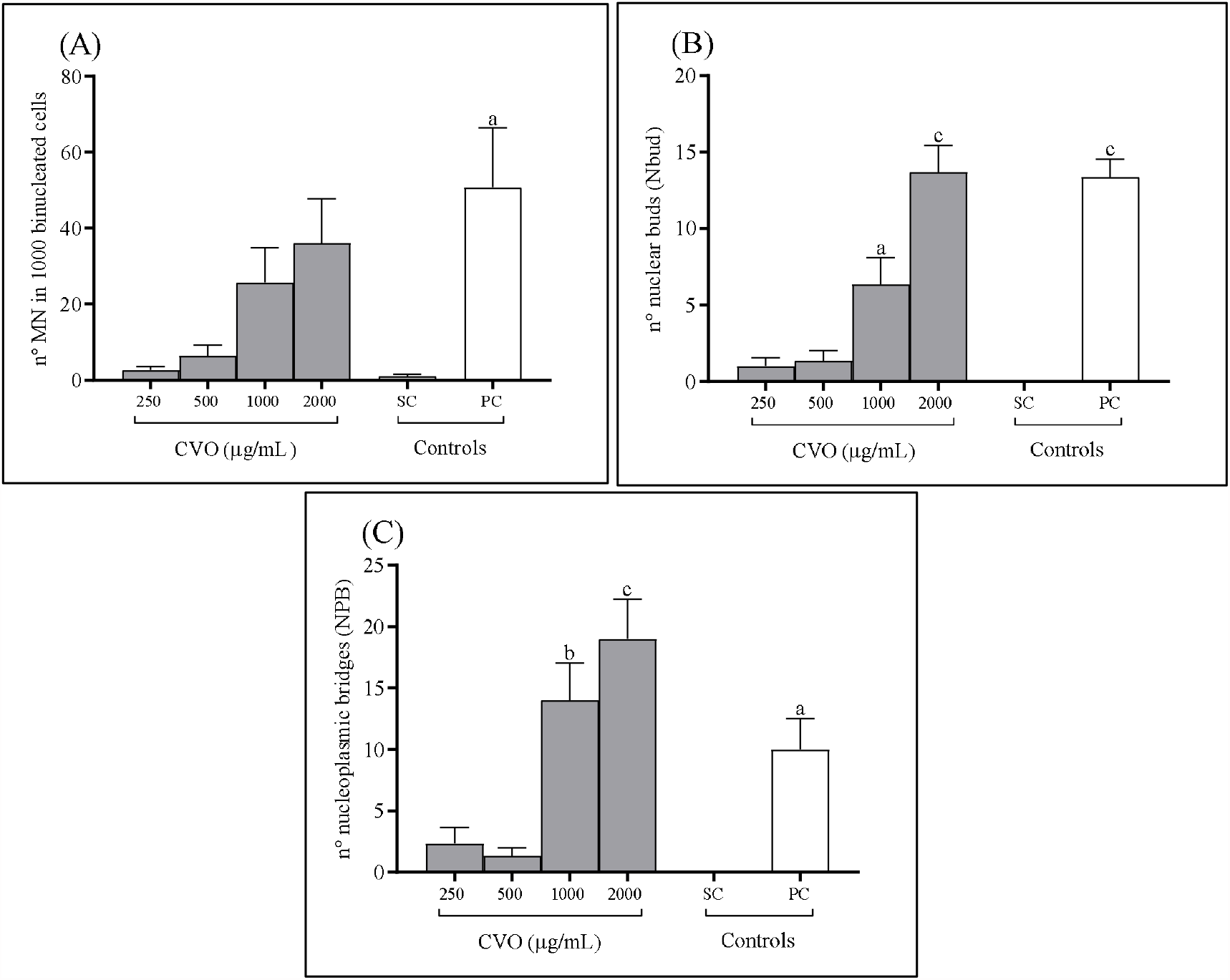
Chromosomal instability in Chinese hamster lung fibroblasts, V79-4 cell line, after treatment with different concentrations of oil extracted from visceral fat of *Caiman yacare* (CVO) for 24 h. (A) Number of micronuclei, (B) nuclear buds, and (C) nucleoplasmic bridges in binucleated V79-4 cells (cytokinesis-block micronucleus assay). Data are presented as mean ± SEM of three independent experiments. One-way ANOVA test, followed by Dunnett’s post-hoc test. Letters above the bars indicate significance; a, p ≤ 0.05; b, p ≤ 0.01; and c, p ≤ 0.001 versus solvent control (CS, 0.08% DMSO). Positive control (PC, doxorubicin 0.025 µg/mL).

## 3. DISCUSSION

The cytotoxic and genotoxic effect of CVO was only observed at the highest concentrations tested in this study. It is worth noting that at lower concentrations, CVO demonstrated antioxidant effects and decreased oxidative DNA damage. These results are similar to those previously described in human retinal pigment epithelial cells (RPE-1) treated with CVO (Azevedo et al., 2020) and murine macrophages treated with *C. siamensis* oil (Ngernjan et al., 2022).

The cytotoxic and cytostatic effects of CVO were accompanied by an increase in ROS production and/or oxidative DNA damage. ROS cause various lesions in DNA, with 8-oxo-guanine, recognized by Fpg glycosylase, being the most common lesion (Cooke et al., 2003). Oxidative DNA damage can activate cell cycle checkpoints, blocking the division progression for damage repair or activation of apoptosis when damages exceed the cell’s processing capacity (Barzilai and Yamamoto, 2004). The cell’s processing capacity is time-dependent, which explains why CVO’s cytotoxic/cytostatic effects are observed as colony formation and cytostasis decreases.

As we described previously, CVO is composed of a nearly equal proportion of saturated (42.95%) and unsaturated (43.74%) fatty acids. Among the saturated fatty acids, palmitic (21.61%) and stearic acids (18.50%) are the most abundant, while oleic acid (32.31%) is the most abundant unsaturated fatty acid. Both saturated fatty acids have been recognized for their *in vitro* cytotoxic effect in T and B cells and in primary cultures of hepatocytes (Lima et al., 2002; Moravcová et al., 2015; Takahashi et al., 2012). In HepG2 cells, palmitic acid increased cytotoxicity and ROS production concomitantly with mitochondrial dysfunction and release of inflammatory markers (Alnahdi et al., 2019), indicating the mechanisms by which this fatty acid can contribute to ROS production.

The CVO also presented mono-(MUFA) and poly-unsaturated fatty acids (PUFA) in its composition, mainly oleic (ômega-9), linoleic, and arachidonic acids (ômega-6). Linoleic and arachidonic acids, when oxidized, generate trans-4-hydroxy-2-nonenal (HNE), which are genotoxic even at low concentrations (Chung et al., 2003; Speit et al., 2004). However, it has been shown that unsaturated fatty acids, such as oleic and linoleic acids, can have beneficial effects at certain concentrations, counteracting the deleterious effects of palmitic acid, decreasing ROS production, DNA damage, and apoptosis (Alnahdi et al., 2019; Beeharry et al., 2003; Moravcová et al., 2015). It is possible that a positive balance between PUFA and saturated fatty acids was achieved at lower, but not at higher concentrations tested in this study, explaining the absence of cytotoxic and genotoxic effects.

As evidenced by the comet assay, treatment with CVO increased DNA breaks. Considering that this effect was also observed at concentrations where there was no increase in oxidative DNA damage, this result suggests that the oil may cause other types of DNA lesions besides oxidative damage. The presence of DNA breaks correlated with higher concentrations of PUFA and saturated fatty acids, but not with an increase in oxidative damage, has been reported previously in lymphocytes *ex vivo* (Thorlaksdottir et al., 2007). However, the mechanism involved in generating this type of lesion remains unclear. Nevertheless, the damage caused at lower concentrations was low and the cells could recover. No increase in the frequency of micronuclei was observed at 250 µg/mL.

The oxidative damage caused at higher concentrations exceeded the cell’s recovery capacity, resulting in increased micronuclei, nuclear buds, and cytoplasmic bridges. These three nuclear anomalies can originate from DNA breaks, which can be caused by the simultaneous repair of proximal oxidative lesions (Fenech et al., 2011; Kisurina-Evgenieva et al., 2016). Oxidative damage can also compromise telomeres, generating chromosomal fusion processes that produce dicentric chromosomes, evidenced as nuclearplasmic bridges (Wang et al., 2010).

Although the higher concentrations tested in this study caused damage that exceeded the cell’s recovery capacity, the concentration of 250 and 500 µg/mL was able to protect the cell from oxidative stress caused by hydrogen peroxide. Interestingly, at this same concentration, CVO showed a lower level of oxidative DNA damage than the solvent control, indicating that it may have some positive influence on lesion removal mechanisms. These results suggest that CVO may have antioxidant or pro-oxidant effects depending on the concentration used.

## 4. MATERIALS AND METHODS

### 4.1 Drugs and reagents

All the following reagents were purchased from Sigma-Aldrich: DCFH-DA (Catalog: D6883, 4091-99-0), Dimethyl sulfoxide (Catalog: D2650, CAS: 67-68-5), Hydrochloride doxorubicin D1515, CAS: 25316-9), N-acetylcysteine (Catalog: A9165-5G, CAS: 616-91-1), Dulbecco′s Modified Eagle′s Medium (DMEM) -Low glucose without phenol red, (Catalog: D2902), Penicillin G sodium salt (Catalog: P3032-10MU, CAS: 69-57-8), Streptomycin sulfate salt (CAS: 3810-74-0) and MTT ([3-(4,5-dimethylthiazol-2-yl)-2,5-diphenyltetrazolium bromide]) (Catalog: M5655-500MG, CAS: 298-93-1). Ciprofloxacin hydrochloride was obtained from Corning (61-277-RG). Hydrogen peroxide was purchased from Supelco (Catalog: HX0636, CAS: 7722-84-1), and formamidopyrimidine [fapy]-DNA glycosylase (Fpg) enzyme (M0240S) from New England BioLabs. DAPI (4’,6-Diamidino-2-henylindole, dihydrochloride) was obtained from Invitrogen (Catalog: D1306, CAS:28718-90-3).

### 4.2 Ethical statement

This study was conducted in accordance with the guidelines of the National Council for the Control of Animal Experimentation (CONCEA), resolution n°. 56, of October 5, 2022, which recognizes alternative methods for the use of animals in research activities in Brazil. We also followed the safety criteria recommended by the National Health Surveillance Agency (ANVISA) and the Organisation for Economic Co-operation and Development (OECD).

### 4.3 Sample preparation

Visceral fats were collected from the *C. yacare* waste produced by zootechnics; the oil was then extracted and the fatty acid profile was characterized as previously described (Azevedo et al., 2020). The mass of crude oil extracted from the visceral fats was determined using an analytical balance (Ohaus Adventure AR2140). The concentrations required for this study were obtained by diluting the crude oil solution (986.6 mg/mL) in dimethyl sulfoxide (DMSO).

### 4.4 Experimental Design

The concentrations selected for the *in vitro* cytotoxicity, intracellular ROS detection, and genotoxicity assays followed the criteria established by the OECD (protocol 487 for the micronucleus assay) (OECD, 2014). However, the solubility of the oil was a limiting factor for obtaining the required concentrations. The lowest amount of solvent required for diluting the stock solution of the oil extracted from visceral fat of *C. yacare* (CVO) was determined to be 1/5 oil and 4/5 Dimethyl sulfoxide -DMSO (v/v).

For the cell viability assay, eight serial concentrations of CVO (15.62– 2000 µg/mL) were selected. After screening using the cell viability assay, four concentrations of CVO (250, 500, 1000, and 2000 µg/mL) were selected for the remaining tests. In all assays, a negative control (NC, culture medium + cells) and solvent control (SC, culture medium + 0.8% DMSO) were used. At this concentration, DMSO was shown to be non-toxic and did not interfere with the results obtained.

For the positive control (PC), two concentrations of doxorubicin were used, 50 mM for the cell viability assay and 0.025 µg/mL for the other tests (comet, micronucleus, and clonogenic survival assays). Hydrogen peroxide (H_2_O_2_) at 1 mM was used as a positive control (PC, H_2_O_2_) and N-acetylcysteine (PC, NAC) at 2 mM was used as a reference control because of its known antioxidant capacity.

### 4.5 Cell Culture

The Chinese hamster lung fibroblast cells of the V79-4 lineage were obtained from the Rio de Janeiro Cell Bank (BCRJ) (code: 0244). The cells underwent karyotyping and were subjected to monthly tests to detect mycoplasma contamination using DAPI staining (Young et al., 2010). The V79-4 lineage were cultured in DMEM culture medium supplemented with 1.5 g/L glucose, 2 mM L-glutamine, and 10% fetal bovine serum (FBS), 0.006% penicillin, and 0.01% streptomycin sulphate. They were maintained in a biochemical oxygen demand (BOD) incubator at 37°C. In all assays, cells were seeded 24 hours prior to the initiation of the treatments.

### 4.6 Cell viability assay

Cell viability analysis was performed using the MTT ([3-(4,5-dimethylthiazol-2-yl)-2,5-diphenyltetrazolium bromide]) assay (Kumar et al., 2018) with modifications. Briefly, a total of 1 × 10^5^ cells were seeded in 96-well plates and treated with different concentrations of CVO and control solutions. After 24 h, the supernatant was removed and the MTT formazan blue precipitate was dissolved in 100 μL of DMSO. The absorbance was measured at 540 nm using spectrophotometry (Thermo Scientific Multiskan EX). Three independent assays and treatments (in triplicate) were conducted. The determination of the percentage of viable cells (%, cell viability) was calculated using the formula:

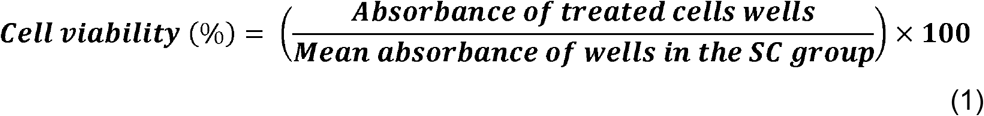

### 4.7 Clonogenic survival assay

The fraction of surviving cells after treatment with CVO was determined using the clonogenic survival assay (Franken et al., 2006) with modifications. A total of 0.14 × 10^6^ cells were seeded in 12-well plates and treated with different concentrations of CVO and control solutions. After 24 h of treatment, the cells were removed with trypsin and 300 viable cells were seeded in a 25 cm^2^ flask per treatment. They were then incubated for 5 d until non-confluent colonies were observed. For colony observation, the cells were fixed with methanol/acetic acid (3:1) and stained with Giemsa. Colonies with at least 50 cells were counted under a binocular stereoscopic microscope (Olympus SZ40). Experiments were performed in triplicate. The number of colonies formed after cell treatment was expressed as a survival fraction (SF), which was calculated from the plating efficiency (PE) value using the following formulas:

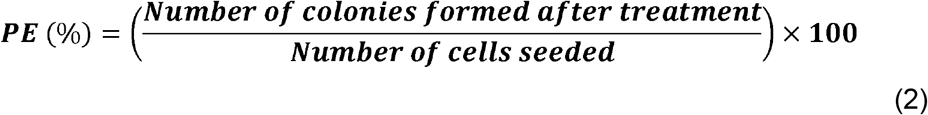

Then, the survival fraction was calculated using the following formula:

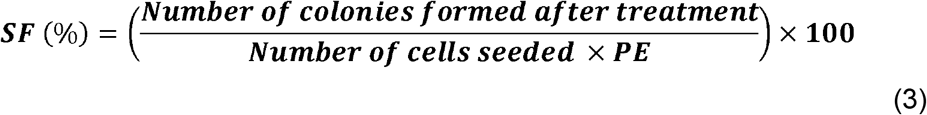

### 4.8 Measurement of intracellular reactive oxygen species (ROS) levels

The intracellular production of reactive oxygen species (ROS) by CVO (pro-oxidant) was analysed by flow cytometry using the fluorescent probe DCFH-DA (LeBel et al., 1992) with modifications. Briefly, cells were seeded at a concentration of 0.14 × 10^6^ cells per well in 12-well plates and treated with different concentrations of CVO and control solutions for 4 h. Then, 500 µL of 5 µM DCFH-DA solution diluted in Hank’s solution was added to each well for 20 min at 37° C in a BOD incubator. Subsequently, excess DCFH-DA was discarded, and cells were washed with Hank’s solution and removed with trypsin. Then, centrifugation was performed for 5 min at 1500 rpm. Next, the supernatant was discarded, and the pellet was suspended in culture medium and Hank’s solution with 2% FBS. Three independent assays and treatments (in duplicate) were conducted. Measurements were recorded using an Accuri C6 plus flow cytometer (BD Biosciences), and event acquisition was performed using the FACSDiva program (BD Biosciences). The analysis of acquired data was performed using Flowjo software (version 10.8.1). ROS production was depicted in histogram graphs, and the mean fluorescence intensity was related to DCFH detection.

To evaluate the protective effect against oxidative stress (antioxidant), cells were pre-treated with the determined concentrations of CVO and control solutions for 3 h. Then, the supernatant was discarded, and cells were washed with Hank’s solution and treated with 1 mM hydrogen peroxide (H_2_O_2_) for 1 h.

The DCFH-DA labelling protocol was followed, and the measurements were recorded as described above.

### 4.9 Comet assay

Single-strand DNA breakage damage was analysed using the standard alkaline comet assay (Singh et al., 1988) with modifications. Briefly, cells were seeded at a concentration of 0.14 × 10^6^ cells per well in 12-well plates and treated with different concentrations of CVO and control solutions for 4 h. Approximately 50 µL of the cell solution was mixed with low-melting-point agarose (0.5%, LM agarose) and placed on slides that were previously prepared with normal-melting-point agarose (1.5%, NM agarose), then incubated at 4 °C for 5 min to polymerize the LM agarose. Lysis, electrophoresis, neutralization, and fixation were then performed as described in the procedure proposed by (de Lucca et al., 2015) with the following modifications: incubation in lysis solution for a minimum of 24 h and incubation in electrophoresis solution for 30 min with a current intensity of 0.86 V/cm (25 V/300 mA) for 30 min at 18° C.

The comet assay and its modified version with the formamidopyrimidine(fapy)-DNA glycosylase (Fpg) enzyme to detect oxidative lesions (8-oxoguanine) caused by ROS were performed as described by (Speit et al., 2004) with modifications. Briefly, cells were analysed under an epifluorescence microscope (Nikon Eclipse Ci ProRes MF) using a 516–560 nm filter, a 590 nm filter barrier (total magnification of 400x), and images were processed using the Lucia Comet Assay software. The percentage of DNA in the tail of cells was used as the standard to measure DNA damage. One hundred nucleoids (50 per slide) were analysed per treatment. Oxidative DNA damage was obtained by calculating the difference between the median values calculated from the percentage of DNA in the tail by treatment in the presence of 1 µg/mL of Fpg enzyme per slide for 30 min and in the absence of the enzyme (Fpg–). Three independent assays and treatments (in duplicate) were conducted.

### 4.10 Cytokinesis-block micronucleus assay (CBMN)

The chromosomal anomalies were analysed in the cytoplasm of interphase cells using the cytokinesis-block micronucleus assay (Fenech, 2007) with modifications. Briefly, approximately 5 × 10^5^ cells were seeded in individual culture flasks (25 cm^2^) for each treatment and treated with cytochalasin B (3 µg/mL) and at the determined concentrations of CVO and control solutions for 24 h (OECD, 2014). Then, the cells were subjected to hypotonic treatment with 2.0 mL of cold sodium citrate solution (1%) and fixed in ice-cold methanol and acetic acid solution (3:1; v/v). Next, the cells were placed onto slides (two slides per treatment) and stained in 3% Giemsa solution diluted in phosphate buffer (pH 6.8) for 5 min. For each treatment, the number of micronuclei (MN), nucleoplasmic bridges (NPB), and nuclear buds (Nbuds) were counted using optical microscopy (at 40x) in 2,000 binucleated cells (1,000 cells per slide). Three independent assays and treatments (in duplicate) were conducted.

The percentage of cytostasis, a cytotoxicity marker, was calculated as follows: 500 cells were counted to determine the proportion of mono-(MN), bi-(BN), and multi-nucleated (MuN) cells, and the cytokinesis-block proliferation index (CBPI) values were determined for each treatment following the formula:

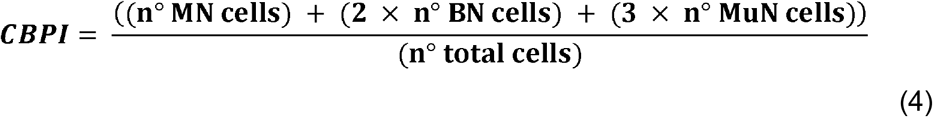

Then, the percentage of cytostasis was calculated using the following formula:

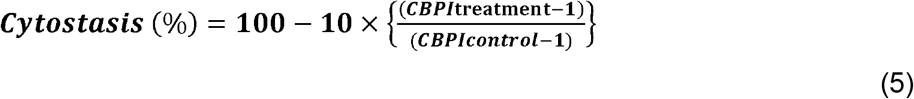

### 4.11 Statistical analyses

The Shapiro-Wilk test was used to verify data distribution. Data presented as the mean ± standard error of mean values (SEM). Means were compared using one-way ANOVA and Dunnett’s post-hoc test. *p* ≤ 0.05 was considered significant for all statistical analyses. Analyses and graphs were created using GraphPad Prism software (version 9.5). No statistical differences were observed between the negative control (NC) and solvent control (SC) in all assays. Thus, considering the presence and non-interference of DMSO in CVO treatments, comparisons were made with respect to SC (MTT, clonogenic survival, comet, and micronucleus assays). For the assay detecting intracellular ROS levels, comparisons were made with respect to positive control (PC, H_2_O_2_) (pro-oxidant activity) and N-acetylcysteine (PC, NAC) (antioxidant activity).

## 5. ACKNOWLEDGMENTS

The authors express their gratitude for the facilities provided by the Laboratory of Scientific Research in Health (LIS), affiliated with the Faculty of Medicine, Federal University of Mato Grosso. *In memory* of our esteemed professor and colleague, Dr. Victor Manuel Aleixo, whose profound dedication to research continues to inspire us. His legacy will always resonate in our work.

## 8. DATA AVAILABILITY

The data generated in this study are available upon reasonable request to the corresponding author.

## REFERENCES

Alnahdi, A., John, A. and Raza, H. (2019). Augmentation of Glucotoxicity, Oxidative Stress, Apoptosis and Mitochondrial Dysfunction in HepG2 Cells by Palmitic Acid. Nutrients 11, 1979. https://doi.org/10.3390/nu11091979.

Alves, R.R.N., Oliveira, T.P.R. and Medeiros, M.F.T. (2017). Trends in Medicinal Uses of Edible Wild Vertebrates in Brazil. Evidence-Based Complement. Altern. Med. 2017, 1–22. https://doi.org/10.1155/2017/4901329.

Azevedo, L.P., dos Santos Ferraz, R.H., de Magalhães, M.R.L., Oliveira, A.P., Cogliati, B., Lemos, L.M.S., Branco, P.C., de Abreu Sousa, D., Cunha da Silva, J.R.M. et al. (2020). Healing potential of Caiman yacare (Daudin, 1802) visceral fat oil. Wound Med. 31, 100195. https://doi.org/10.1016/j.wndm.2020.100195.

Barzilai, A. and Yamamoto, K.-I. (2004). DNA damage responses to oxidative stress. DNA Repair (Amst). 3, 1109–1115. https://doi.org/10.1016/j.dnarep.2004.03.002.

Beeharry, N., Lowe, J.E., Hernandez, A.R., Chambers, J.A., Fucassi, F., Cragg, P.J., Green, M.H. and Green, I.C. (2003). Linoleic acid and antioxidants protect against DNA damage and apoptosis induced by palmitic acid. Mutat. Res. Mol. Mech. Mutagen. 530, 27–33. https://doi.org/10.1016/S0027-5107(03)00134-9.

Campos, Z., Llobet, A., Magnusson, W.E. and Piña, C. (2020). Caiman yacare. The IUCN Red List of Threatened Species 2020. https://dx.doi.org/10.2305/IUCN.UK.2020-3.RLTS.T46586A3009881.

Chung, F.L., Pan, J., Choudhury, S., Roy, R., Hu, W. and Tang, M.S. (2003). Formation of trans-4-hydroxy-2-nonenal- and other enal-derived cyclic DNA adducts from ω-3 and ω-6 polyunsaturated fatty acids and their roles in DNA repair and human p53 gene mutation. Mutat. Res. - Fundam. Mol. Mech. Mutagen. 531, 25–36. https://doi.org/10.1016/j.mrfmmm.2003.07.001.

Cooke, M.S., Evans, M.D., Dizdaroglu. and M., Lunec, J. (2003). Oxidative DNA damage: mechanisms, mutation, and disease. FASEB J. 17, 1195–1214. https://doi.org/10.1096/fj.02-0752rev.

de Lucca, R.M.R., Batista Júnior, J., Fontes, C.J.F., de Oliveira Bahia and M., Bassi-Branco, C.L. (2015). Genotoxic effects of the antimalarial drug lumefantrine in human lymphocytes in vitro and computational prediction of the mechanism associated with its interaction with DNA. Environ. Mol. Mutagen. 56, 556–562. https://doi.org/10.1002/em.21942.

Fenech, M. (2007). Cytokinesis-block micronucleus cytome assay. Nat. Protoc. 2, 1084–1104. https://doi.org/10.1038/nprot.2007.77.

Fenech, M., Kirsch-Volders, M., Natarajan, A.T., Surralles, J., Crott, J.W., Parry, J., Norppa, H., Eastmond, D.A. Tucker. et al. (2011). Molecular mechanisms of micronucleus, nucleoplasmic bridge and nuclear bud formation in mammalian and human cells. Mutagenesis 26, 125–132. https://doi.org/10.1093/mutage/geq052.

Abrão, C.F., de Oliveira, D.R., Passos, P., Freitas, C.V.R.P, Santana, A.F., Lopes da Rocha, M., Ribeiro da Silva, A.J. and Wanderley Tinoco, L. (2021). Zootherapeutic practices in the Amazon Region: chemical and pharmacological studies of Green-anaconda fat (Eunectes murinus) and alternatives for species conservation. Ethnobiol. Conserv. 10, 1–31. https://doi.org/10.15451/ec2021-02-10.15-1-27.

Franken, N.A.P., Rodermond, H.M., Stap, J., Haveman, J. and van Bree, C. 2006. Clonogenic assay of cells in vitro. Nat. Protoc. 1, 2315–2319. https://doi.org/10.1038/nprot.2006.339.

Kisurina-Evgenieva, O.P., Sutiagina, O.I. and Onishchenko, G.E. (2016). Biogenesis of micronuclei. Biochem. 81, 453–464. https://doi.org/10.1134/S0006297916050035.

Kramer, J.A., Sagartz, J.E. and Morris, D.L. (2007). The application of discovery toxicology and pathology towards the design of safer pharmaceutical lead candidates. Nat. Rev. Drug Discov. 6, 636–649. https://doi.org/10.1038/nrd2378.

Kumar, P., Nagarajan, A. and Uchil, P.D. (2018). Analysis of Cell Viability by the MTT Assay. Cold Spring Harb. Protoc. 2018, 469–471. https://doi.org/10.1101/pdb.prot095505.

Kumar, S., Dhir, A., Talwar, S., Chakraborty, D. and Kaur, P. (2021). What drives brand love for natural products? The moderating role of household size. J. Retail. Consum. Serv. 58, 102329. https://doi.org/10.1016/j.jretconser.2020.102329.

LeBel, C.P., Ischiropoulos, H. and Bondy, S.C. (1992). Evaluation of the Probe 2′,7′-Dichlorofluorescin as an Indicator of Reactive Oxygen Species Formation and Oxidative Stress. Chem. Res. Toxicol. 5, 227–231. https://doi.org/10.1021/tx00026a012

Li, H.-L., Liu, X.-T., Huang, S.-M., Xiong, Y.-X., Zhang, Z.-R., Zheng, Y.-H., Chen, Q.-X. and Chen, Q.-H. (2021). Repair function of essential oil from Crocodylus Siamensis (Schneider, 1801) on the burn wound healing via up-regulated growth factor expression and anti-inflammatory effect. J. Ethnopharmacol. 264, 113286. https://doi.org/10.1016/j.jep.2020.113286.

Lima, T.M., Kanunfre, C.C., Pompéia, C., Verlengia, R. and Curi, R. (2002). Ranking the toxicity of fatty acids on Jurkat and Raji cells by flow cytometric analysis. Toxicol. Vitr. 16, 741–747. https://doi.org/10.1016/S0887-2333(02)00095-4.

Mishra, B., Akhila MV, A., Thomas, A., Benny, B. and Assainar, H. (2020). Formulated Therapeutic Products of Animal Fats and Oils: Future Prospects of Zootherapy. Int. J. Pharm. Investig. 10, 112–116. https://doi.org/10.5530/ijpi.2020.2.20.

Moravcová, A., Červinková, Z., KuČera, O., Mezera, V., Rychtrmoc, D. and Lotková, H. (2015). The Effect of Oleic and Palmitic Acid on Induction of Steatosis and Cytotoxicity on Rat Hepatocytes in Primary Culture. Physiol. Res. 64, S627–S636. https://doi.org/10.33549/physiolres.933224.

Ngernjan, M., Ontawong, A., Lailerd, N., Mengamphan, K., Sarapirom, S. and Amornlerdpison, D. (2022). Crocodile Oil Modulates Inflammation and Immune Responses in LPS-Stimulated RAW 264.7 Macrophages. Molecules 27, 3784. https://doi.org/10.3390/molecules27123784.

Organization for Economic Co-operation and Development (2014). Test No. 487: In Vitro Mammalian Cell Micronucleus Test. Paris, France: OECD Publishing https://doi.org/10.1787/9789264224292-en.

Praduptong, A., Siruntawineti, J., Chaeychomsri, S., Srimangkornkaew, P. and Chaeychomsri, W. (2018). Acute oral toxicity testing of Siamese crocodile (Crocodylus siamensis) oil in Wistar rats. Biosci. Discov. 9, 409–415.

Romanelli, P. and Schmidt, J. (2003). Estudo do aproveitamento das vísceras do jacaré do pantanal (Caiman yacare) em farinha de carne. Ciência Tecnol. Aliment. 23, 131–139.

Shim-Prydon, G. and Camacho-Barreto, H (2007). New animal products: new uses and markets for by-products and coproducts of crocodile, emu, goat, kangaroo and rabbit.

Singh, N.P., McCoy, M.T., Tice, R.R., Schneider, E.L. (1988). A simple technique for quantitation of low levels of DNA damage in individual cells. Exp. Cell Res. 175, 184–191. https://doi.org/10.1016/0014-4827(88)90265-0.

Speit, G., Schütz, P., Bonzheim, I., Trenz, K., Hoffmann, H. (2004). Sensitivity of the FPG protein towards alkylation damage in the comet assay. Toxicol. Lett. 146, 151–158. https://doi.org/10.1016/j.toxlet.2003.09.010.

Takahashi, H.K., Cambiaghi, T.D., Luchessi, A.D., Hirabara, S.M., Vinolo, M.A.R., Newsholme, P., Curi, R. (2012). Activation of survival and apoptotic signaling pathways in lymphocytes exposed to palmitic acid. J. Cell. Physiol. 227, 339–350. https://doi.org/10.1002/jcp.22740.

Thorlaksdottir, A.Y., Jonsson, J.J., Tryggvadottir, L., Skuladottir, G. V., Petursdottir, A.L., Ogmundsdottir, H.M., Eyfjord, J.E. and Hardardottir, I. (2007). Positive Association Between DNA Strand Breaks in Peripheral Blood Mononuclear Cells and Polyunsaturated Fatty Acids in Red Blood Cells From Women. Nutr. Cancer 59, 21–28. https://doi.org/10.1080/01635580701365092.

Vicente-Neto, J., Bressan, M.C., Faria, P.B., Vieira, J.O., Cardoso, M. das G., Glória, M.B. de A. and da Gama, L.T. (2010). Fatty acid profiles in meat from Caiman yacare (Caiman crocodilus yacare) raised in the wild or in captivity. Meat Sci. 85, 752–758. https://doi.org/10.1016/j.meatsci.2010.03.036.

Wang, Z., Rhee, D.B., Lu, J., Bohr, C.T., Zhou, F., Vallabhaneni, H., de Souza-Pinto, N.C. and Liu, Y. (2010). Characterization of oxidative guanine damage and repair in mammalian telomeres. PLoS Genet. 6, 28. https://doi.org/10.1371/journal.pgen.1000951

Young, L., Sung, J. and Masters, J.R. (2010). Detection of mycoplasma in cell cultures. Nat. Protoc. 5, 929–934. https://doi.org/10.1038/nprot.2010.43.

